# An integrative framework for circular RNA quantitative trait locus discovery with application in human T cells

**DOI:** 10.1101/2023.03.22.533756

**Authors:** Dat Thanh Nguyen

**Affiliations:** Centre for Integrative Genetics, Faculty of Biosciences, Norwegian University of Life Sciences, Ås, Norway

**Keywords:** circular RNA, circQLT, eQTL, collocation, GWAS

## Abstract

Molecular quantitative trait locus (QTL) mapping of genetic variants with intermediate molecular phenotypes has proven to be a powerful approach for prioritizing genetic regulatory variants and causal genes identified by Genome-wide association studies (GWAS). Recently, this success has been extended to circular RNA (circRNA), a potential group of RNAs that can serve as markers for the diagnosis, prognosis, or therapeutic targets of cancer, cardiovascular, and autoimmune diseases. However, the detection of circRNA QTL (circQTL) currently is heavily reliant on a single circRNA detection algorithm for circRNA annotation and quantification which implies limitations in both sensitivity and specificity. In this study, we show that circQTL results produced by different circRNA calling tools are extremely divergent, making difficulties in interpretation. To resolve this issue, we develop an integrative method for circQTL mapping and implement it as an automated, reproducible, and scalable, and easy-to-use framework based on Nextflow, named cscQTL. Compared to the existing approach, the new method effectively identify circQTLs with an increase of 20-100% circQTLs detected and recovered all circQTLs that are highly supported by the single method approach. We apply the new method to a dataset of human T cells and discover genetic variants that control the expression of 55 circRNAs. By collocation analysis, we further identify circBACH2 and circYY1AP1 as potential candidates for immune disease regulation. cscQTL is freely available at: https://github.com/datngu/cscQTL.

## 1 Introduction

In order to understand how genetic variations contribute to the variation of phenotypes, genome-wide association studies (GWAS) are often employed to identify such relations. Over nearly two decades of accumulation, GWAS have successfully detected thousands of DNA variants associated with human complex and disease traits^[1]^. Furthermore, we have learned that complex traits are mostly driven by non-coding variants with small effect sizes^[2,3]^. Although the number of detected trait-associated variants continues to grow, understanding the underlying molecular mechanisms of most GWAS loci remains a challenge due to the non-coding nature of 90% GWAS loci^[4,5]^.

Among the possible approaches to tackle this challenge, molecular quantitative trait locus (QTL) mapping of genetic variants with intermediate molecular phenotypes, such as gene expression (eQTLs) and splicing (sQTLs), has proven to be a powerful tool to prioritize genetic regulatory variants and causal genes. Multiple studies have shown that GWAS hits are enriched in significant eQTLs/sQTLs loci and regulatory elements, suggesting gene regulation mechanisms of trait-associated DNA variants^[6,7,8,9,10,11]^. Moreover, the success of the QTL approach has recently expanded to include the discovery of genetic variants controlling RNA editing^[12]^.

Circular RNA (circRNA) is a relatively young class of RNA molecules characterized by covalently closed-loop structures without a 5’ cap or a 3’ poly (A) tail, formed by back-splicing events during RNA splicing processes^[13,14,15]^. To date, over a million circRNAs have been identified in humans and other vertebrate species^[16]^. Multiple studies showed that circRNAs exhibit unique expression patterns in tissues and developmental stages and are more stable than other RNA types^[17,18,19]^. Functional studies suggested that circRNAs play critical roles in various cellular processes and disease pathogenesis, including acting as microRNA sponges^[20,21]^, regulating premRNA splicing^[22]^, and modulating innate immunity^[23]^. Indeed, a few pioneer circQTL studies shed the light on the impact of genetic variants on circRNA expressions to regulatory mechanisms underlying human complex diseases^[24,25,26,27]^.

It is different from linear RNA with high-quality annotations available for direct quantification. CircRNAs typically lack full-length transcript annotations and must be identified and quantified primarily based on the detection of reads containing a back-splicing junction (BSJ), where the end of an exon joins to the start of itself or of another exon from the same gene. In addition, circRNAs are known to express differently across tissues^[17,18,19]^ making circRNA identification becomes necessary in circQTL studies.

While eQTL is well established with multiple computational pipelines and standardized protocols exist^[9,10,28,29]^, circQTL mapping is still in its early stages with no comprehensive computational pipeline available. Current circQTL studies are commonly implemented by simply taking the output of a single circRNA detection algorithm, and directly using the BSJ counts as expression levels for QTL mapping^[24,25,26,27]^. This type of circQTL mapping is hereafter referred to as the single method approach. Although it is easy to use, utilizing only one circRNA calling method implies certain limitations. Firstly, circRNA detection still suffers from a certain amount of false positives regardless of the efforts of state-of-the-art circRNA calling methods^[30,31,32]^. Secondly, circRNA detection exhibits little agreement between calling tools that implies the potential divergence results in circQTL downstream analyses^[33,34,35,36]^. One potential solution to improve the specificity and sensitivity of circRNA detection is combining outputs of several circRNA detection algorithms^[36]^. However, this approach is only straightforward for annotation studies such as circRNA database construction^[16]^. For quantitative studies such as circQTL, it would arise another technical challenge in quantification integration since each circRNA calling algorithm has its own output of expression levels.

Here, we aim to establish the first comprehensive computational pipeline for circQTL analysis. We show that using a single circRNA detection method approach indeed produces highly divergent QTL results between circRNA calling algorithms. To address this challenge, we develop an integrative method called cscQTL to systematical combine circRNA output from different tools for circQTL analysis by a re-quantification approach. Compared to the single method circQTL approach, cscQTL identifies more circQTLs and provides more coherence results. We implement cscQTL as an automated, reproducible, and scalable framework based on Nextflow^[37]^. By applying cscQTL, we find genetic variants controlling expressions of 55 circRNAs in human T cells and identify circBACH2 and circYY1AP1 as potential circRNAs for immune disease regulation by collocation analyses.

## 2 Results

### 2.1 Challenges in circQTL analysis

To assess the impact of different circRNA calling algorithms on circQTL mapping results, we evaluate three commonly used algorithms: Circall^[32]^, CIRI2^[31]^, and CIRCexplorer2^[30]^ using a publicly available dataset of 40 individuals^[38]^. These algorithms are chosen based on their reported high accuracy and efficiency in previous studies^[32,34,35]^. We apply each algorithm with its default settings for circRNA calling, using the same genome annotation. We then perform uniform data reprocessing and circQTL mapping, as described in the Methods section.

The number of circRNAs remaining for QTL testing is 5586, 6778, and 3336 for Circall, CIRI2, and CIRCexplorer2, respectively. To confirm the accuracy of the called circRNAs, we compare them against the most comprehensive public catalog of CircAtlas^[16]^. The majority of the called circRNAs by all three methods are found in CircAtlas, with ratios of 97.15%, 97.42%, and 99.79% for Circall, CIRCexplorer2, and CIRI2, respectively (Table 1 and Fig 1 A). We observe a low degree of overlap between the calling algorithms, which is consistent with previous studies^[34,35]^. Specifically, out of the 7224 unique circRNAs called by the three algorithms, 41.89% (3026/7224) are detected by all three methods, and 75.44% (5450/7224) are supported by at least two methods (Fig 1 B).

**Table 1.** The number of circRNA used for QTL testing with reference to CircAtlas and the number of eCircRNAs detected by three circRNA calling algorithms Circall, CIRI2, CIRCexplorer2, and their consensus statistics.

| Methods | Detected circRNAs |  | Detected eCircRNA |
| --- | --- | --- | --- |
|  | All | CircAtlas |  |
| Circall | 5586 | 5427 | 46 |
| CIRI2 | 6778 | 6764 | 41 |
| CIRCexplorer2 | 3336 | 3250 | 15 |
| Total unique circRNA/eCircRNA | 7224 | 7015 | 71 |
| Supported by at least 2 methods | 5450 | 5402 | 24 |
| Supported by all 3 methods | 3026 | 3024 | 7 |

**Fig 1.**
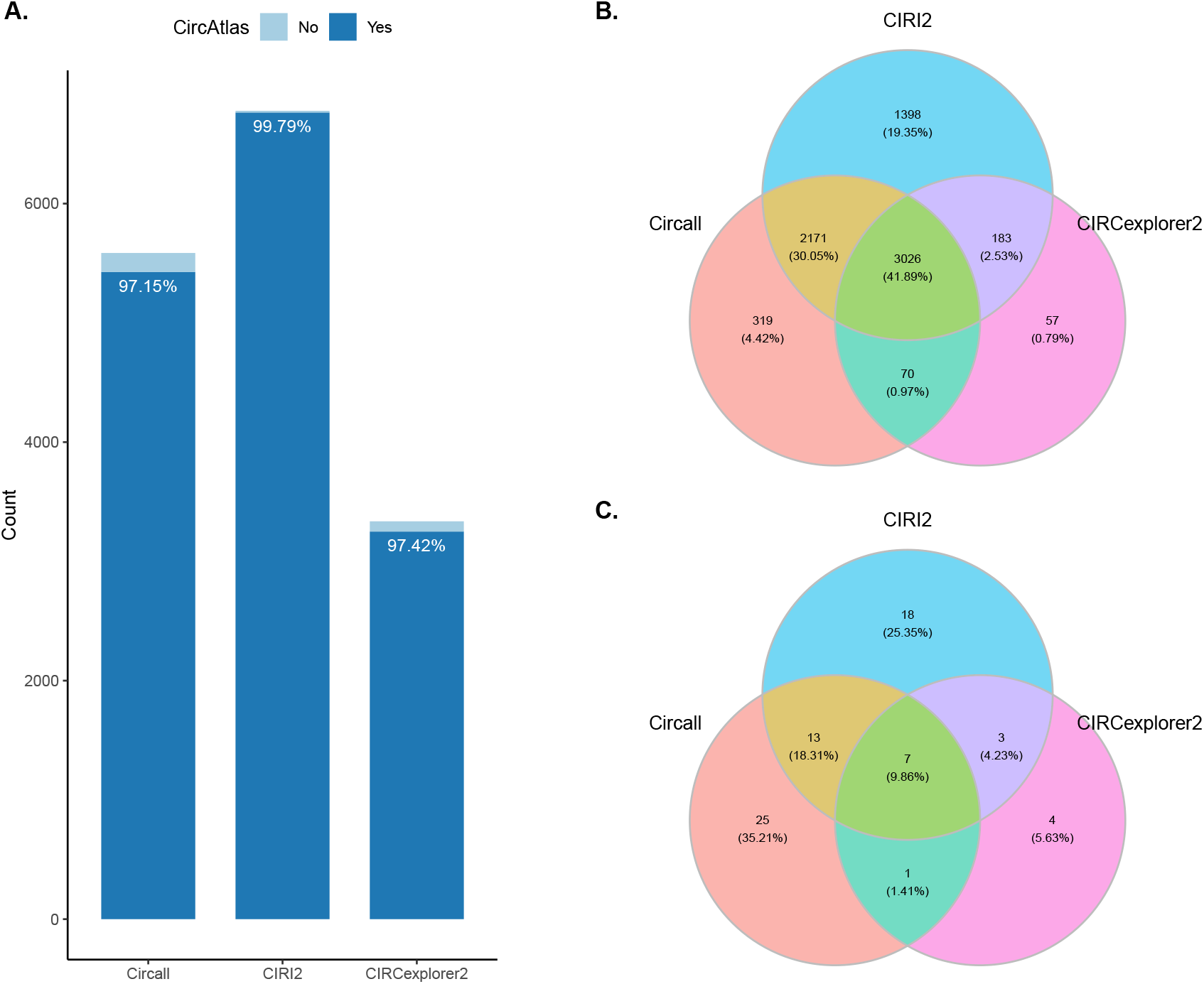
Challenges in circQTL analysis. (A) The number of circRNAs used for QTL testing with reference to CircAtlas. (B) and (C) Venn diagrams showing the overlapping of QTL-tested circRNAs and eCircRNAs identified by three different circRNA calling algorithms.

Regarding the QTL mapping results, a total of 71 distinct eCircRNAs (circRNAs whose expression levels associate with at least one genetic variant) are detected under Storey’s q-value < 0.05 procedure^[39]^. Among these, 46 were identified by Circall, 41 by CIRI2, and 15 by CIRCexplorer2. Less than 10% (7/71) of the detected eCircRNAs are supported by all three algorithms, and approximately one-third (24/71) is supported by at least two methods, as shown in Table 1 and Fig 1 C. These results indicate that the choice of circRNA detection tools affects both the number and content of eCircRNAs detected. Although circRNA detection is a well-known challenging task, with results varying by the choice of algorithm^[33,34,35]^, the consensus of QTL mapping results is even more divergent indicating limitations in both sensitivity and specificity of single method circQTL mapping approach. Furthermore, this inconsistent results can make interpretation intractable, limit the transferability and reproducibility of the analysis, and hence, require a more efficient method to address these issues.

### 2.2 cscQTL: integrative circular RNA quantitative trait locus discovery

#### 2.2.1 Overview of the cscQTL pipeline

Motivated by previous studies demonstrating that combining multiple circRNA calling tools and re-quantification can improve consistency in circRNA calling and downstream differential expression analyses^[36,40]^, we propose a novel integrative framework called cscQTL (consensus-based circRNA QTL mapping) to address these challenges. Firstly, cscQTL minimizes divergence among circRNA calling algorithms by combining circRNA inputs from three high-accuracy circRNA identification algorithms. Secondly, cscQTL implements re-mapping and quantification procedures to provide accurate quantification of circRNAs. Finally, cscQTL is implemented using Nextflow^[37]^, which enables reproducible and scalable circQTL analyses in an automatic and user-friendly manner.

An overview of cscQTL is presented in Fig 2 A. First, RNA-seq (ribo-minus) data is used for circRNA identification by Circall, CIRI2, and CIRCexplorer2, with specific aligners allocated based on the authors’ suggested parameters. The detected circRNA candidates are harmonized to obtain a similar format before consensus-based filtering to obtain high-quality BSJ sites. In this implementation, circRNA candidates with >= 2 BSJ reads in at least one sample are kept before consensus filtering with different cutoffs of 1, 2, and 3 supporting methods. To accurately quantify the expression level of circRNAs and filter false-positive BSJ reads, reads that are fully mapped to the linear transcripts are discarded, and quantification is performed by counting the number of fragments mapped to pseudo circRNA references generated by concatenating 149 bases of the upstream sequences of the end positions with 149 bases of the downstream sequences of the start positions (Fig 2 B). Quasi-mapping is used for these alignment steps to ensure computational efficiency^[41]^. Since only highly confident circRNAs are considered in the quantification, a loose filtering criterion is applied, i.e., filtering is applied with only one condition that the shorter piece of the read must cover the junction break point with at least 7 bases. After obtaining the counting matrix, the subsequent steps, such as population filtering, z-score scaling, quantile-quantile (Q-Q) normalization, covariate analyses, and QTL mapping, proceed similarly to other methods before collocation testing, as described in detail in the Methods section.

**Fig 2.**
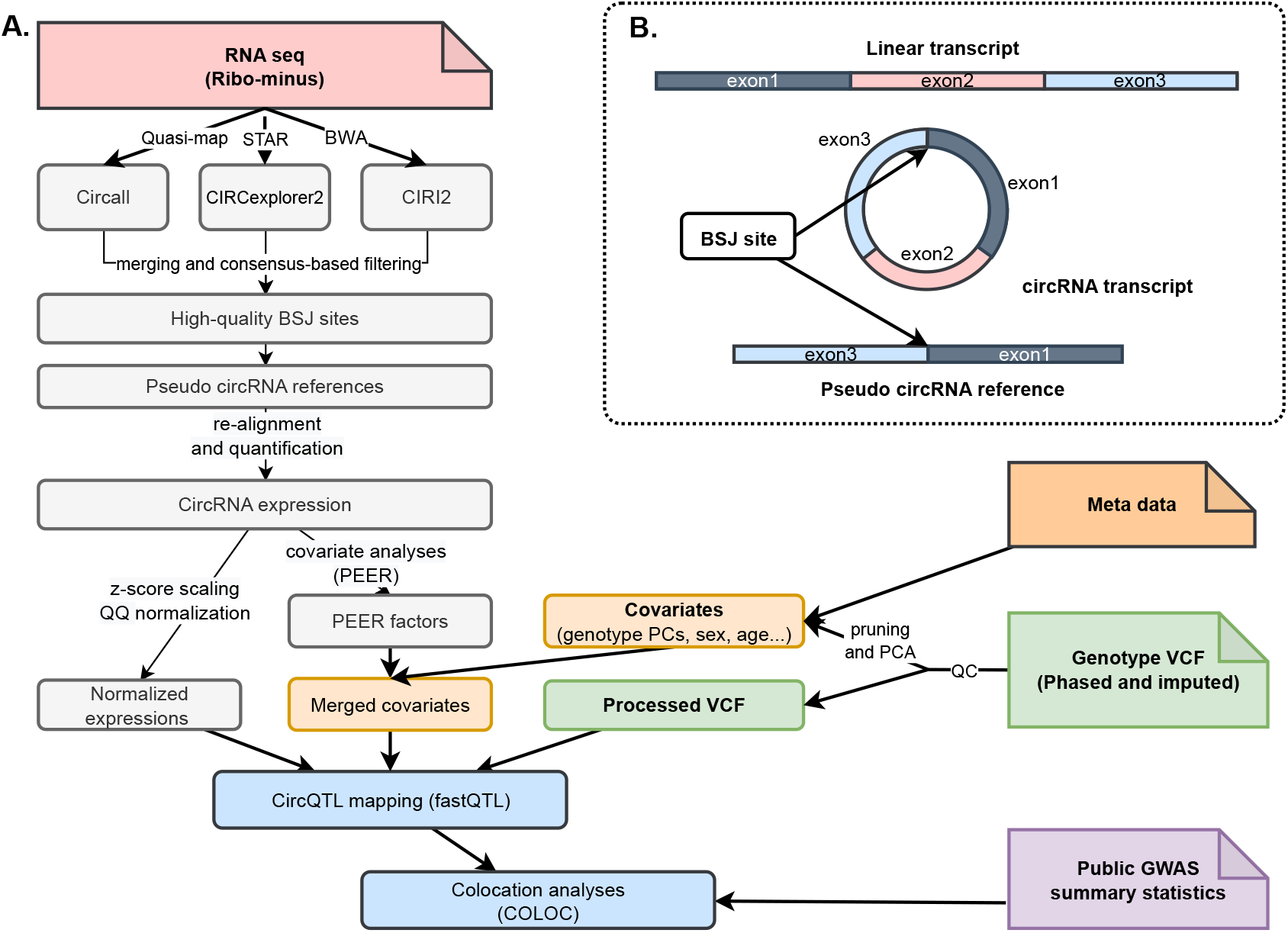
Overview of the cscQTL pipeline. A) Workflow of circQTL mapping. CircRNAs are first identified by Circall, CIRCexplorer2, and CIRI2. Then, cscQTL applies consensus-based filtering to obtain high-quality circRNA candidates before quantifying by re-alignment of RNA-seq reads against the pseudo circRNA references; circRNA expressions then go through scaling, quantile-quantile normalization before CircQTL mapping and collocation analyses. B) The construction of pseudo circRNA references. Based on circRNA annotations detected by circRNA calling algorithms, a pseudo circRNA reference is generated by joining 149 bases of the upstream sequences of the end positions with 149 bases of the downstream sequences of the start positions. It is noted the pseudo circRNA reference is not present in the corresponding linear form.

#### 2.2.2 Comparison of cscQTL against standard circQTL mapping approaches

To illustrate the efficiency of cscQTL, we apply the pipeline with different consensus cutoffs of 1, 2, and 3 denoted as cscQTL_1, cscQTL_2, and cscQTL_3 respectively, and compare them against the single method QTL approach that performs circQTL mapping by directly use circRNA quantification from a single circRNA calling method as implemented with Circall, CIRI2, and CIRCexplorer2. Since we do not know the ground truth of genetic variants controlling circRNA expressions. We consider the result of the single method circQTL mapping approach as the baseline for evaluating the concordance and the number of eCircQTL called.

In this experiment, we also check the contents of tested circRNAs to circAtlas to illustrate the validity of the analyses (Fig S.1 A). Regarding circQTL mapping, eCircRNAs detected are completely overlapped (Fig S.1 B) and and the numbers are decreased by the levels of stringency as expected. At the loosest setting (consensus cutoffs of 1), cscQTL detected a total of 94 unique eCircRNAs. The corresponding numbers are 61, and 55 for the setting of 2, and 3 that cscQTL considers circRNA candidates supported by at least 2 or 3 methods by either Circall, CIRI2, and CIRCexplorer2 for re-quantification. Compared to the single method approach, cscQTL identifies approximately 20-100% more eCircRNAs (depending on consensus settings) than the most effective single method approach which is Circall with 46 eCircRNA identified as shown in 3 A. Importantly, cscQTL results are highly concordant with the single method circQTL mapping. For instance, 40 out of 71 eCircQTL identified all three single methods are recalled by cscQTL_3. The corresponding number of cscQTL_2 and cscQTL_1 are 41 and 51 out of 71. Furthermore, all 24 highly confident eCircRNAs that are identified by at least 2 single methods are showing up in all consensus settings. Overall, these results indicate the robustness of cscQTL (Fig 3 B, and Fig S.1 C, D).

**Fig 3.**
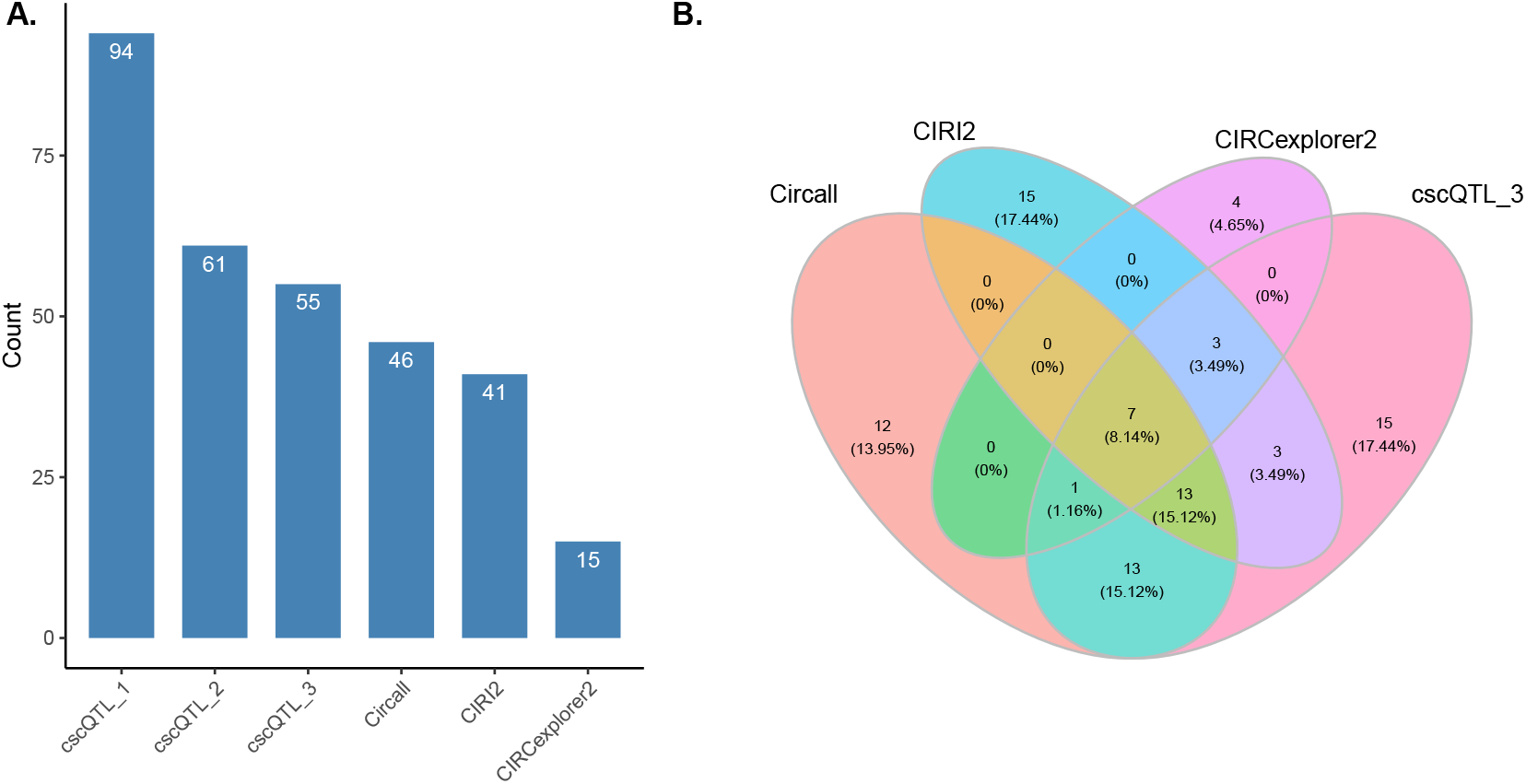
Comparison of circQTL calling pipelines. (A) The number of eCircRNAs detected by Circall, CIRCexplorer2, CIRI, and cscQTL (tested circRNAs are supported by at least 1, 2, and 3 circRNA detection algorithms corresponding to cscQTL_1, cscQTL_2, and cscQTL_3). (B) Venn diagrams showing the overlap of eCircQTLs identified by Circall, CIRCexplorer2, CIRI, and cscQTL_3.

### 2.3 Genetic regulation of circRNA in human T cell

#### 2.3.1 Characterization of circRNA-associated genetic variants

As an application of cscQTL, we set the consensus setting to 3, the most stringent setup, for further analyses. We tested approximately 6.1 million SNPs and 6080 circRNAs for cis-QTL associations within 1MB of BSJ boundaries. As shown in Figure 4A, the Q-Q plot of the genome-wide distribution of the test statistic against the expected null distribution is extremely higher than the diagonal line in the right tail, indicating no evidence for systematic spurious associations. By applying permutation tests implemented in FastQTL and Storey’s q-value < 0.05 multiple testing correction procedure for the number of circRNAs tested, we identify 55 eCircRNAs that belong to 45 distinct host genes according to Ensembl v106 annotation. To identify the full list of significant variant-circRNA associations, we use the top nominal p-value of the circRNA closest to the 0.05 FDR threshold as a genome-wide threshold. In this way, we identify a total of 5269 pairs of significant variant-circRNA associations. We define genetic variants with at least a significant in this context as circSNPs. These circSNPs are then further annotated with the Ensembl Variant Effect Predictor (VEP)^[42]^. The most dominant variant classes were intronic, upstream gene, and intergenic variants, accounting for 73.37%, 6.21%, and 6.06% respectively, while other classes account for the remaining proportion, as shown in Figure 4B.

**Fig 4.**
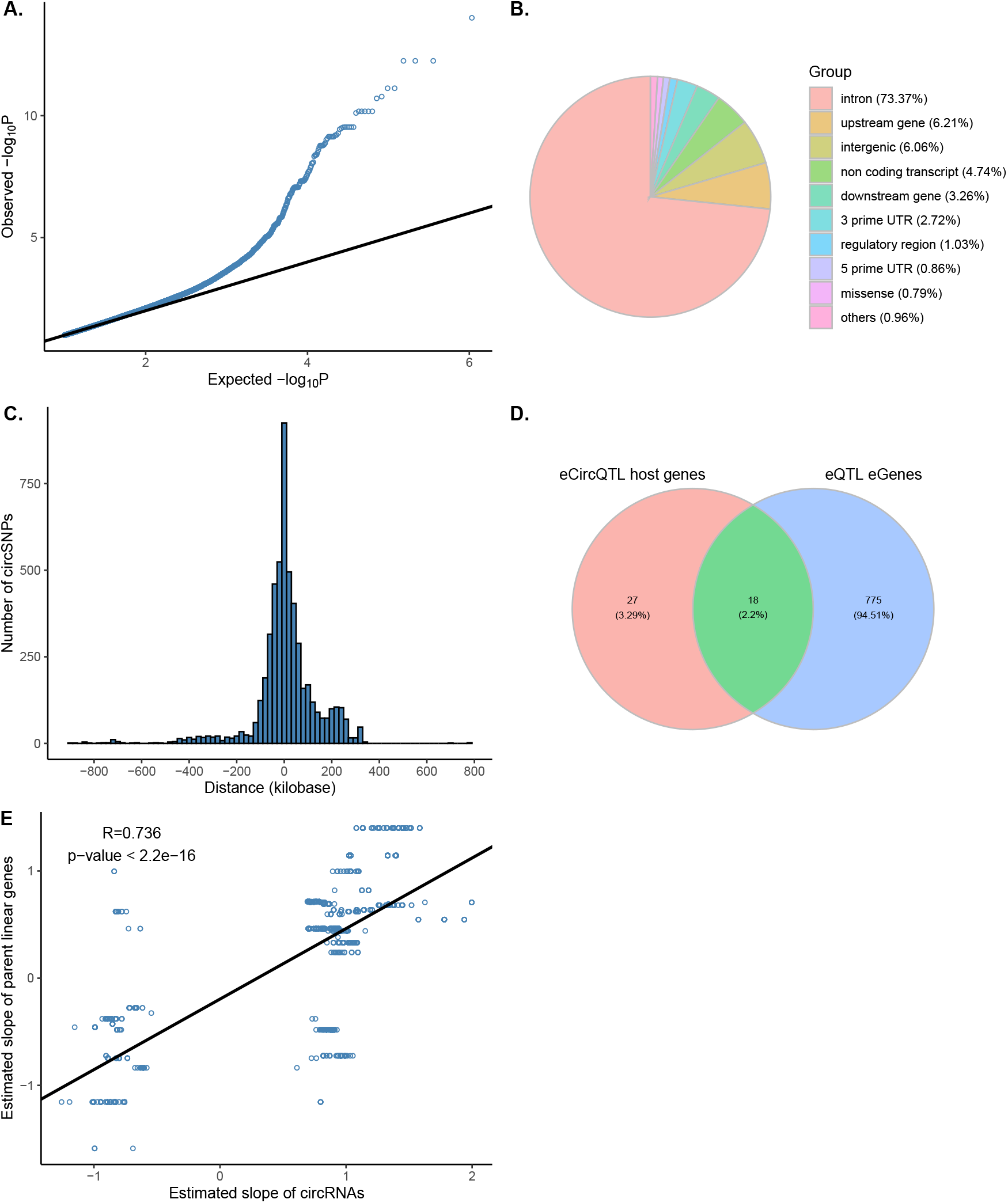
Genetic regulation of circRNA expression in human T cells. (A) Q–Q plot distribution of all nominal p-values. (B) Predictive effects of circSNPs by Variant Effect Predictor. (C) Distance distribution of circSNPs from the BSJ sites. (D) Overlapping between circQTL host genes and gene-based eQTLs. (E) Pearson correlation and its significant test of circSNPs’ effect sizes and the effect size of their corresponding parent mRNA genes.

Regarding the distance distribution relative to the tested circRNAs, circSNPs are mostly located near the BSJ, with rare cases located beyond 500kb from BSJs, as shown in Figure 4C. To understand the relationship between circQTLs and their host gene eQTLs, we further perform a traditional gene-based eQTL analysis in the same dataset (details are described in Methods). We observe that around 35% (16/45) of eCircRNA host genes are also significant in gene-based tests, regardless of the larger number of eGenes identified with 793 distinct Ensembl gene IDs, as shown in Figure 4D indicating the distinction between the two classes of RNA. We finally compare the effect sizes of the two QTLs by searching for matched pairs of variants and circRNA host genes. We observe a strong correlation (Pearson correlation R=0.736, p-value < 2.2e-16) of effect sizes between gene-based eQTL and circQTLs, as shown in Figure 4E.

#### 2.3.2 Collocation between circQTLs and immune disease GWAS loci

To investigate the possible relation between circQTLs and immune GWAS loci, we perform collocation tests using COLOC^[43]^. Specifically, we obtain GWAS summary statistics of Crohn’s disease (CD), Inflammatory bowel disease (IBD)^[44]^, and Type 1 diabetes (T1D)^[45]^ from the GWAS catalog webpage^[46]^. We consider a PP.H4 >= 0.5 as the collocation threshold and visualize the collocation using LocusCompare^[47]^. Overall, two out of 55 circRNAs exhibit collocations with immune disease GWAS loci including circBACH2 (6:90206569:90271941 - ENSG00000112182) and circYY1AP1 (1:155676548:155679512 - ENSG00000163374). circBACH2 is collocated with all tested traits including T1D, CD, and IBD with probabilities of 0.89, 0.91, and 0.91, respectively (Fig 5 A, S.2, and S.3). Regarding circYY1AP1, it is collocated to CD with a probability of 0.68 and to IBD with a probability of 0.61 (Fig S.4 and S.5). Interestingly, BACH2 is a known risk gene for T1D^[48]^. We further investigate the effect sizes of the top circSNPs (rs60066732) in circQTL and gene-level eQTL. As shown in Fig 5 B and C, the SNP rs60066732 are negatively correlated with both types of transcripts, but the effect size, as well as the p-value of the circRNA transcript, are much more extreme and significant than its host gene, indicating the potential role of circRNA in T1D risk.

**Fig 5.**
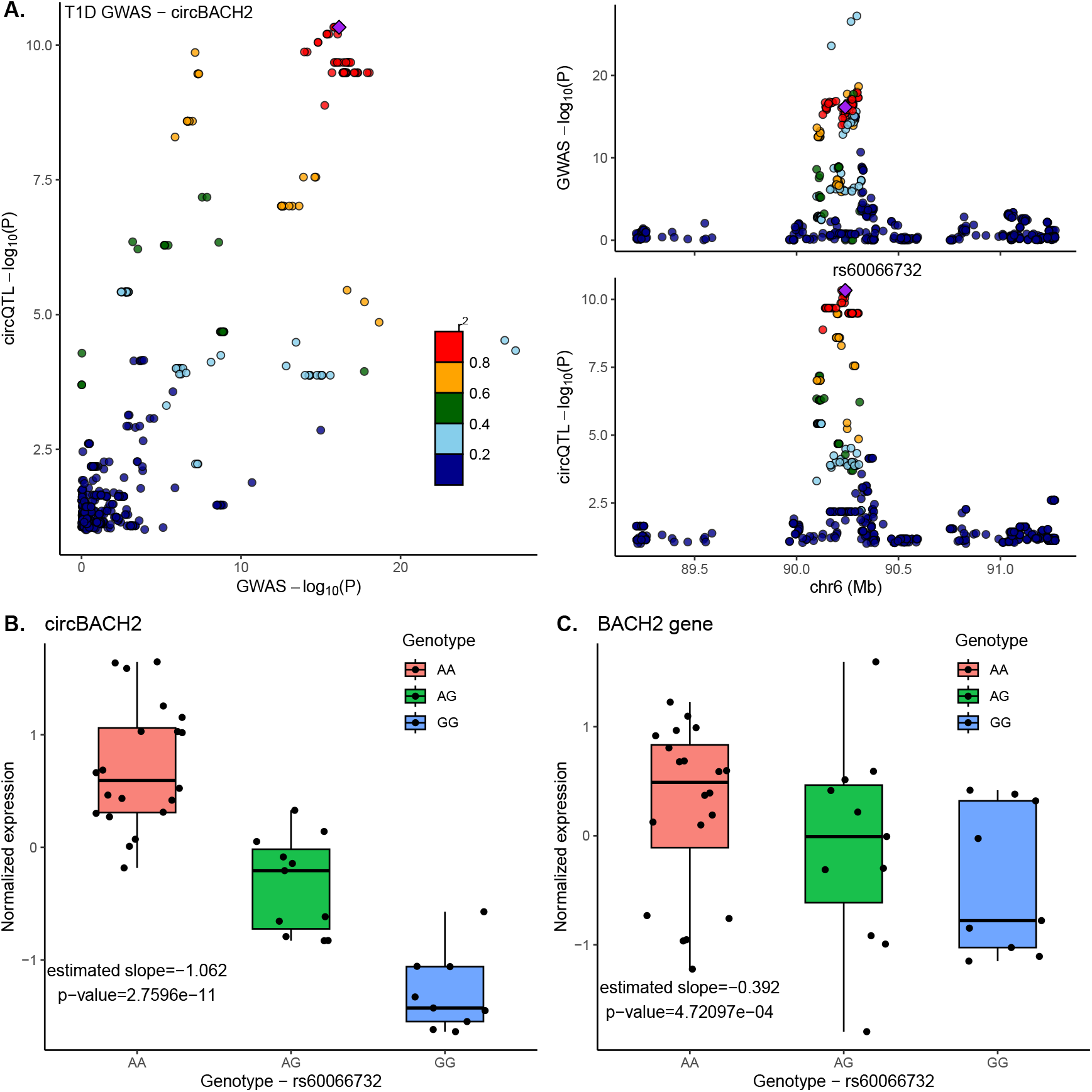
A T1D GWAS locus associated with circBACH2 (6:90206569:90271941) expression. (A) circBACH2 collocated with T1D GWAS. The top SNP rs60066732 is highlighted with a purple diamond. (B) and (C) Associations of circBACH2 and its linear parent gene (BACH2) to the top SNP with the risk allele G.

## 3 Discussion

CircRNA detection is known as a challenging task. Deploying a single algorithm in a circQTL study indeed exhibits highly divergent results suggesting limitations in both sensitivity and specificity of the approach. Combining several algorithms in circRNA detection has been proposed and implemented in database construction^[16,36]^ as an efficient solution for these issues. In this study, we extend this idea to circQTL analysis by developing a novel computational framework called cscQTL to unify outputs of multiple circRNA calling algorithms for circQLT mapping. By using consensus-based filtering together with the re-quantification procedure, cscQTL provides a coherent interpretation of circQTLs while it is still able to take advantage of combing multiple circRNA calling algorithms. Compared to the single-method circQTL mapping approach, cscQTL recalls all highly confident circQTLs (identified by at least two single methods). The method further identifies more circQTLs circQTLs than any single-method circQTL mapping even with the most stringent setting (considers only circRNA candidates identified in all three algorithms), indicating its reliability and robustness. While the high concordant with single method circQTL mapping can be explained by the consensus-based filter, the efficiency can be explained by the accurate re-quantification procedure of cscQTL. In contrast to cscQTL which focuses only on the quantification of high-quality circRNA candidates identified by combining several algorithms. CircRNA detection algorithms are commonly designed for de-novo identification in small sample size experiments with various statistical and rule-based BSJ read filtering implemented^[49]^. Although sophisticated filters are useful in reducing possible false positive circRNAs in sample-based circRNA discovery, they may introduce unexpected noise in population-based downstream analyses that directly use BSJ counting as expression levels.

Performing circQTL analysis is a fairly complicated task that often employs multiple bioinformatics frameworks and statistical tests, limiting their transferability and replicability. Given that current circQTL studies make use of highly customized pipelines. For instance, Ahmed et al’s study^[24]^ employed their in-house pipeline for circRNA calling, combined with Matrix eQTL^[50]^, and eigenMT^[51]^ for multiple testing correction while Aherrahrou et al made use of CIRI2^[31]^, tensorQTL^[52]^, and q-value correction^[39]^ for the corresponding tasks. With the increasing attention on circRNAs, the bioinformatics community would benefit from a unified but open-source and portable circQTL workflow. By taking the advantage of Nextflow, we implement cscQTL as an easy-to-use pipeline and made it freely available to the public. With inputs being a directory of RNA-seq data, a genome reference, GWAS summary statistic data, and a metadata input file, cscQTL automatically performs circRNA identification, filtering, re-quantification, QTL mapping, and collocation analyses. As the potential role of circRNA in human health and disease is becoming more appreciated^[53,54,55,56]^, we believe that our proposed method will facilitate the discovery of circQTLs in the near future.

Overall, this study provides insights into the impact of circRNA calling tools on circQTL results and presents an improved method for circQTL mapping that can potentially enhance the understanding of the regulatory mechanisms underlying circRNAs in human diseases

## 4 Methods

### 4.1 Datasets

A publicly available RNA-depleted RNA sequencing dataset consisting of 40 individuals, including CD4+ T naive cells, are obtained from Synapse (https://doi.org/10.7303/syn22250947). Briefly, purified and enriched CD4+ T cell samples were processed using the Illumina TruSeq Stranded Total RNA Kit with Ribo-Zero Gold, and subsequently, sequenced on the Illumina HiSeq 2000 platform, as previously described^[38]^, to produce 100bp paired-end (PE) reads. Corresponding whole-genome sequencing genotypes of matched individuals are also downloaded from Figshare (https://doi.org/10.6084/m9.figshare.12646238.v5), as provided in the same publication^[38]^.

### 4.2 Circular RNA identification

The downloaded ribo-minus RNA-seq reads are used for circRNA calling using three different tools: Circall v1.0.1^[32]^, CIRI v2.0.6^[31]^, and CIRCexplorer2^[30]^. The reference genome, transcriptome, and genome annotation of hg38 version 106 are obtained from the Ensembl website (http://ftp.ensembl.org/pub/release-106/). All tools are run with their default suggested parameters and designed alignment algorithms. For instance, Circall v1.0.1 is executed with the option “-td TRUE” and its built-in quasi-mapping^[41]^; CIRI2 is applied with BWAMEM^[57]^ parameter “–T 19”; and CIRCexplorer2 is deployed with the STAR aligner^[58]^ with options “–chimSegmentMin 10 –chimOutType Junctions,” as suggested in a benchmarking study^[35]^.

### 4.3 Data processing and QTL mapping

To ensure a fair evaluation between circRNA calling tools used in the study, all detected circRNAs are uniformly processed. Initially, the detected circRNAs from each tool are merged. To filter out lowly expressed circRNAs, we apply a population-scale filter to keep only those with supporting BSJ counts greater than or equal to 2 and detected in at least 30% of the 40 samples. The BSJ counts are then normalized by library sizes to obtain circRNA count per million units (CPM). The resulting expression matrices are then scaled and quantile-quantile normalized to fit a normal distribution as implemented in standard eQTL workflows^[11,59]^.

Regarding the genotyping data, variants with Hardy-Weinberg Equilibrium (HWE) p-values below 10-6 and minor allele frequency below 5% are removed using PLINK v1.9^[60]^. Additionally, genetic principal components (PCs) are calculated after pruning using the options “–indep-pairwise 200 50 0.25” to exclude potential confounding effects due to population genetic structure.

QTL mappings are then performed using FastQTL v2.184^[61]^. For each test feature (circRNA/mRNA), the adaptive permutation mode is used with the setting with options “–permute 1000 10000” within 1MB window “–window 1e6”. The obtained FastQTL beta distribution approximated empirical p-values are then used to calculate trait-level q values^[39]^ to account for multiple testing. A false discovery rate (FDR) < 0.05 is then applied to identify eCircRNA .i.e. circRNA with at least one significant circQTL. For covariate inclusion in the linear regression, sex, age, four genotype PCs, and a number of PEER factors (between from 1 to 20)^[62]^ are included to optimize the number of significant eCircRNA. The optimized PEER factor is then used for nominal QTL mapping to obtain nominal p-values for downstream analyses. Finally, the detected circQTLs are tested for collocation with GWAS loci using COLOC^[43]^. In this implementation, we consider several immune disease GWAS summary statistics, including Crohn’s disease, inflammatory bowel disease^[44]^, and type 1 diabetes^[45]^. Significant GWAS loci are tested with all significant eCircRNAs with a threshold of PP.H4 >= 0.5.

### 4.4 Gene-based eQTL mapping

Regarding gene-based eQTL testing, we follow a similar procedure as for circQTL with some modifications in quantification. Specifically, we use the pseudo-alignment mode of Salmon v1.3.0^[63]^ to directly quantify RNA-seq data, using the same genome annotation (hg38 version 106 obtained from Ensembl). We then aggregate transcript-level quantification of mRNAs in transcript per million (TPM) units to the gene level using the “tximport” package^[64]^. We also apply the population-scale filter to remove genes expressed in less than 30% of the 40 samples before performing data scaling and quantile-quantile normalization. We then perform eQTL mapping with FastQTL^[61]^ and account for multiple testing using the adaptive permutation scheme and q-value procedure^[39]^, as described in the previous section.

## Ethics approval and consent to participate

The study does not generate any new dataset. Ethics approval and consent to participate were applied according to the corresponding original works.

## Competing interests

The authors declare NO competing interests.

## Availability of data and materials

sccQTL is available at: https://github.com/datngu/cscQTL. Results data support the finding and source codes to generate figures for this study are available at: https://github.com/datngu/cscQTL_paper.

## Author contributions

DTN conceived the study, collected datasets, developed the software, analyzed data, interpreted results, and wrote the manuscript.

## Acknowledgments

The author especially thanks Thong H. Nguyen and Lien T.K. Tran for supporting in maintaining the computing unit. I also thank my advisors including Lars Grønvold, Simen Rød Sandve, and Sigbjørn Lien for the free to finish this work. I special thanks to Nghia Vu and Lars Grønvold for carefully reading the manuscript and for insightful suggestions.

## Funding

This work is supported by NMBU PhD Research Fellowship.

## Notes

https://github.com/datngu/cscQTL

https://github.com/datngu/cscQTL_paper

https://doi.org/10.7303/syn22250947

https://doi.org/10.6084/m9.figshare.12646238.v5

### Competing Interest Statement

The authors have declared no competing interest.

