## Supplementary figures for "An integrative framework for circular RNA quantitative trait locus discovery with application in human T cells"

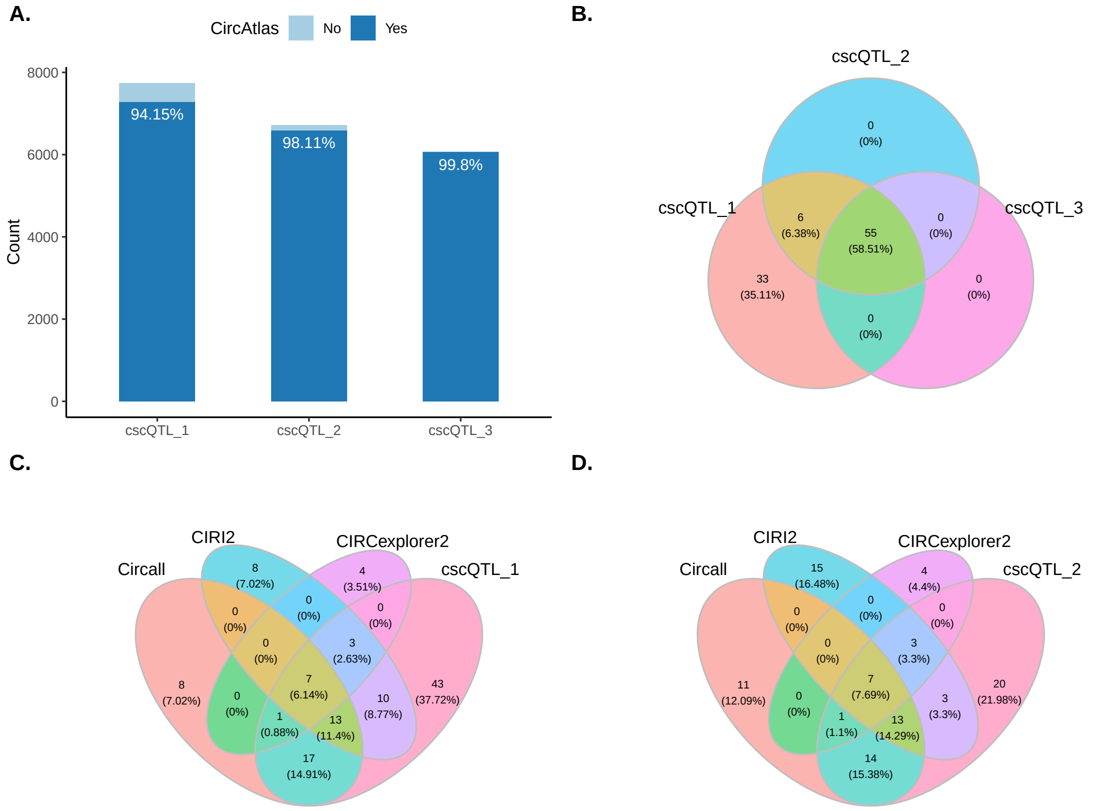


**Fig S. 1** (A) The number of circRNAs used for QTL testing with referencing to CircAtlas of cscQTL with various cutoffs. (B) enn diagrams showing the overlapping of cscQTL_1, cscQTL_2, and cscQTL_3. (C) Venn diagrams showing the overlap of eCircQTLs identified by Circall, CIRCexplorer2, CIRI, and cscQTL_1. (C) Venn diagrams showing the overlap of eCircQTLs identified by Circall, CIRCexplorer2, CIRI, and cscQTL_3

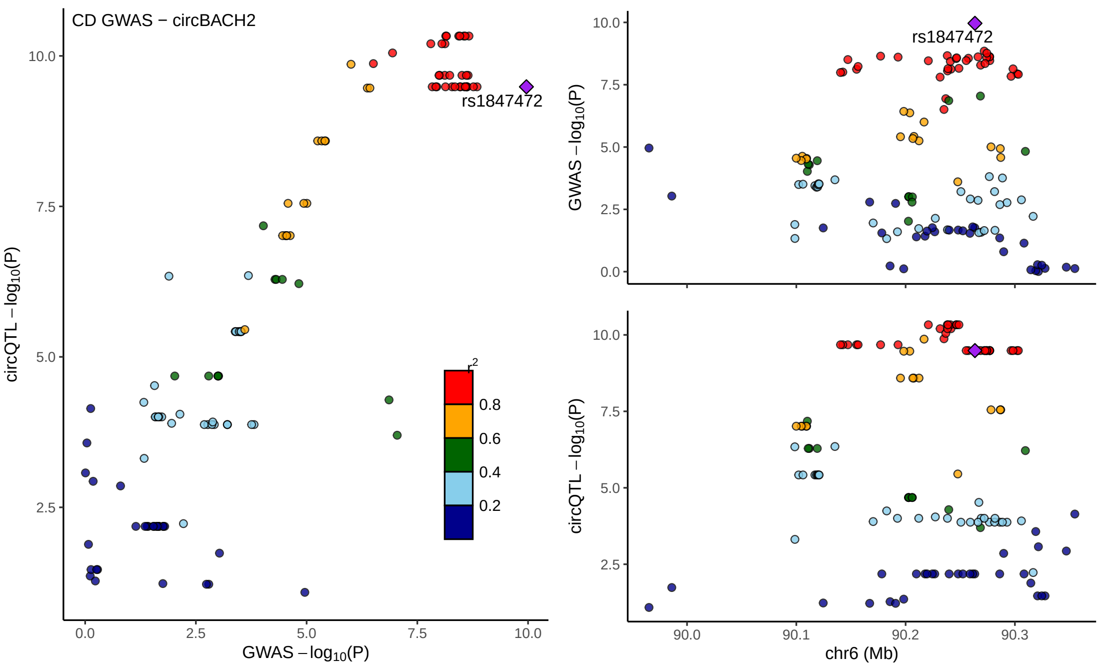


**Fig S. 2** A CD GWAS locus associated with circBACH2 (6:90206569:90271941)

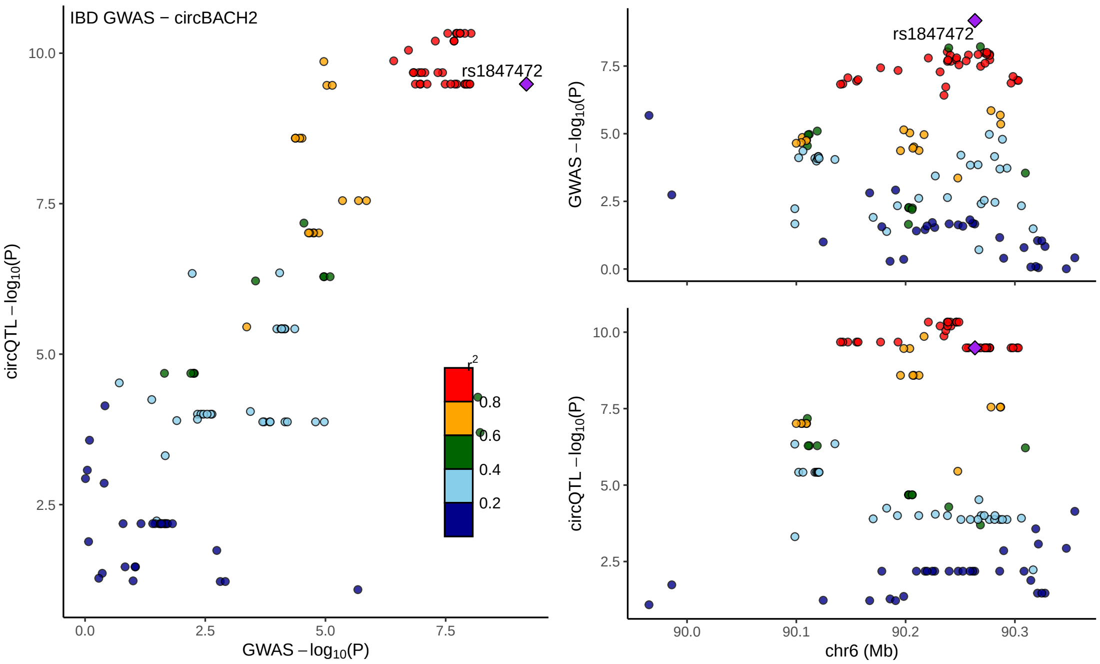


**Fig S. 3** An IBD GWAS locus associated with circBACH2 (6:90206569:90271941)

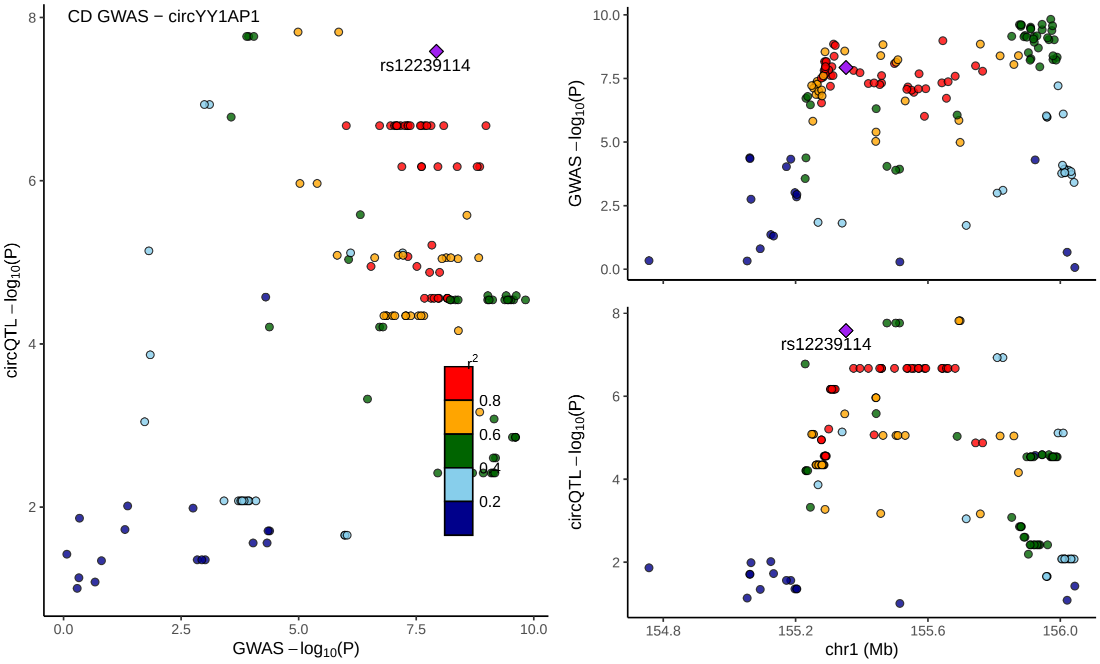


**Fig S. 4** An CD GWAS locus associated with circYY1AP1 (1:155676548:155679512 )

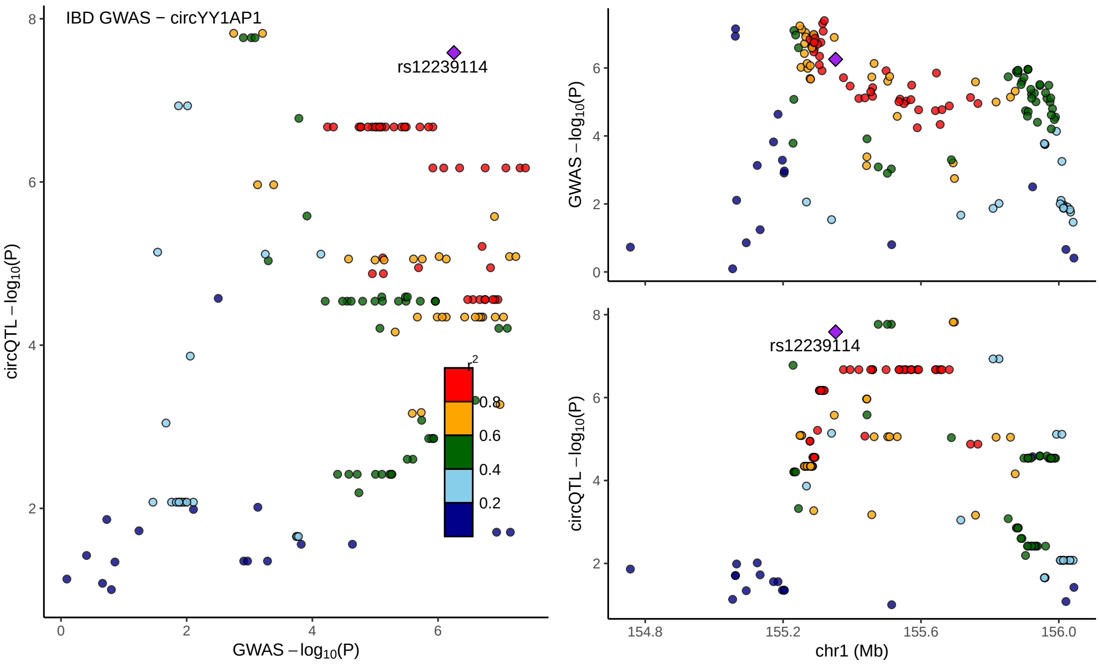


**Fig S. 5** An IBD GWAS locus associated with circYY1AP1 (1:155676548:155679512)
